# Ancient climate changes and relaxed selection shape cave colonization in North American cavefishes

**DOI:** 10.1101/2024.11.20.624529

**Authors:** Pamela B. Hart, Melissa Rincon-Sandoval, Fernando Melendez-Vazquez, Jonathan W. Armbruster, Emily M. Troyer, Orran M. Bierstein, Brendan J. Gough, Ricardo Betancur-R, Matthew L. Niemiller, Dahiana Arcila

**Author notes:** these authors contributed equally (Pamela B. Hart, Dahiana Arcila).

## Abstract

Extreme environments serve as natural laboratories for studying evolutionary processes, with caves offering replicated instances of independent colonisations. The timing, mode, and genetic underpinnings underlying cave-obligate organismal evolution remains enigmatic. We integrate phylogenomics, fossils, paleoclimatic modeling, and newly sequenced genomes to elucidate the evolutionary history and adaptive processes of cave colonisation in the study group, the North American Amblyopsidae fishes. Amblyopsid fishes present a unique system for investigating cave evolution, encompassing surface, facultative cave-dwelling, and cave-obligate (troglomorphic) species. Using 1,105 exon markers and total-evidence dating, we reconstructed a robust phylogeny that supports the nested position of eyed, facultative cave-dwelling species within blind cavefishes. We identified three independent cave colonisations, dated to the Early Miocene (18.5 Mya), Late Miocene (10.0 Mya), and Pliocene (3.0 Mya). Evolutionary model testing supported a climate-relict hypothesis, suggesting that global cooling trends since the Early-Middle Eocene may have influenced cave colonisation. Comparative genomic analyses of 487 candidate genes revealed both relaxed and intensified selection on troglomorphy-related loci. We found more loci under relaxed selection, supporting neutral mutation as a significant mechanism in cave-obligate evolution. Our findings provide empirical support for climate-driven cave colonisation and offer insights into the complex interplay of selective pressures in extreme environments.

## Introduction

Life finds a way: despite the harsh physiological demands of environments such as sulfidic springs or 400°C deep-sea hydrothermal vents, organisms form communities in these habitats. Understanding the origin and maintenance of traits that allow for survival in extreme conditions is of central interest to biologists because these traits are tightly linked to organismal fitness and ecological resilience (1, 2). Beyond just convergent phenotypes (e.g., temperature-stable enzymes in microorganisms, antifreeze glycoproteins in ice fishes) associated with extreme environments in disparate taxa (3–5), recent studies have demonstrated that genetic signatures of convergence may be more common across the genome of different species in the Tree of Life than previously considered (6–8).

Caves exemplify environments that impose harsh physiological demands on their resident organisms. Cave-obligate organisms have evolved to life in complete darkness and often limited energy and nutrient sources, which has led to the evolution of a convergent series of morphological, physiological, and behavioural traits (i.e., troglomorphy) associated with living in subterranean environments. Troglomorphies include the loss/reduction of pigmentation and eyes and enhancement of non-visual sensory perception across invertebrate and vertebrate taxa (9–13). Fishes are the most species-rich vertebrate group to colonise and permanently reside in caves with over 200 species across ten fish orders (14–16). Most exhibit convergent troglomorphic traits that have become ubiquitous with cave-obligate evolution (i.e., reduction of eye structures, vision, and pigmentation) (12, 13, 15).

Several hypotheses have been proposed to explain the colonization of caves and other subterranean habitats from surface environments. Leading hypotheses for subterranean speciation invoke evolutionary processes (modes of speciation; e.g., allopatry vs. parapatry and sympatry) and environmental factors (abiotic vs. biotic factors) (17–20). Paleoclimatic shifts have been hypothesised as a potential factor associated with cave colonization (i.e., caves as refugia or climate relicts) (21–25). The climate-relict hypothesis suggests that organisms pre-adapted to low-light and low-resource habitats, initially inhabited both surface and subterranean habitats with gene flow. After significant climatic events, surface populations went extinct, while cave-dwelling populations persisted and adapted (21). Alternatively, colonization might occur via incidental entry into subterranean habitats, as hypothesised for species like the Mexican blind cave tetra (26, 27) and Hawaiian planthoppers (28). Biotic factors, such as the absence of predators and the lower interspecific competition, may have provided a suitable environment for further adaptation and the evolution of troglomorphic characters (29). While subterranean adaptive radiations are rare (30–33), examples like the *Niphargus* amphipods in European cave systems highlight the potential for diversification in these ecosystems (34).

The family Amblyopsidae, part of the entirely North American order Percopsiformes, offer an excellent system for studying cave adaptation and evolution. The members exhibit different stages of cave-obligate evolution: surface (one species), facultative cave-dwelling (surface-cave; two species), and cave-obligate (six species). Amblyopsidae, along with its sister families Percopsidae (two extant species) and Aphredoderidae (five extant species) (35, 36), inhabits a diverse range of habitats across three major karst regions: the Appalachians, Interior Low Plateau, and Ozarks (17, 37–41). These karst landscapes, formed of soluble carbonate rock, such as limestones and dolomites, are rich in cave systems, yet stratigraphy varies between the karst regions. This geological diversity provides a unique backdrop for the evolution of cave-dwelling species.

The evolutionary history of Amblyopsidae has remained unclear due to discrepancies between morphological and molecular data (17, 38, 42–44), particularly concerning the facultative cave-dwelling genus *Forbesichthys*, which possesses small functional eyes, retains pigmentation, and occupies both surface and subterranean habitats. Morphological data suggest that cave-obligate fishes are monophyletic, with facultative cave-dwelling fishes as their sister group (45). However, molecular data indicate a paraphyletic relationship, positioning facultative cavefishes as the sister group to the cave-obligate genus *Amblyopsis* (26, 46–48). This suggests either a single eye-loss event followed by a reversal or multiple independent eye-loss events (46, 47). These findings make Amblyopsidae an ideal model for investigating the interplay between habitat, adaptation, and evolutionary history in cave-dwelling species.

Here, we present comprehensive phylogenomic and comparative genomic analyses of the North American cavefishes (Percopsiformes: Amblyopsidae). Our study aims to: (1) reconstruct the evolutionary history and timing of cave colonization events in this group; (2) test hypotheses regarding the factors associated with cave colonization, particularly the role of paleoclimate; and (3) identify genomic signatures of adaptation to cave environments. Using newly generated whole-genome sequencing data for 13 species, including multiple individuals per species across the surface-to-cave habitat continuum, we provide insights into the genomic basis of troglomorphy and the evolutionary processes shaping cave adaptation. We also employ a Bayesian phylogenomic framework that incorporates both fossils (four species) and extant percopsiform species, using the fossilised birth death model to estimate divergence times. This integrative approach, combining phylogenomics, paleoclimate modeling, and selection analysis, provides a framework for exploring the relationships between environmental change, genomic evolution, and phenotypic adaptation in other systems.

## Results and Discussion

### Phylogenomic analyses and divergence time estimation

Our time-calibrated percopsiform phylogeny, based on extensive taxon sampling incorporating both fossil and extant percopsiforms identified three shifts from surface to cave habitats. These shifts occurred approximately 18.5, 10.0, and 3.0 million years ago. We found support for a topology that included eyed, facultative cave-dwelling fishes nested within blind, and cave-obligate fishes. Although it is unlikely that the transition from non-cave to cave environments occurred precisely at the moment a cavefish lineage began diverging in the phylogenetic tree, we used the confidence intervals of the nodes for branching cavefish lineages as a proxy for when cave colonization may have begun. Previous studies support at minimum three colonization events in the amblyopsid cavefishes, one at each clade (*Tr. rosae*, *Amblyopsis* spp., and *Typhlichthys* spp. + *Speoplatyrhinus poulsoni*) (17, 38, 43); therefore, we estimated cave colonization for all three of these nodes in the phylogeny. Additional evidence suggests multiple independent colonization events within the cave-adapted genera in the family (17, 37, 43).

Our topology is congruent with the most recent molecular studies of the Percopsiformes and Amblyopsidae (17, 38, 43), resolving the eyed, facultative cave-dwelling *Forbesichthys* species as the sister group to the Northern blind cave-obligate fishes (*Amblyopsis* spp.) (Fig. 1, Supp Mat S2 Fig. S1). The *Forbesichthys* + *Amblyopsis* clade is the sister group to all other cave-obligate fishes. The surface Swampfish *Chologaster cornuta* is the first-branching taxa of the Amblyopsidae (Fig 1). The first (oldest) shift from surface to cave environments occurred 18.5 Mya (11–29.8 Mya, 95% HPD; Fig. 1A). The second cave colonization took place 10.0 Mya (4.1–17.3 Mya, 95% HPD; Fig. 1B). The most recent surface-to-cave change occurred 3.0 Mya (0.3–7.1 Mya, 95% HPD; Fig. 1C). Our divergence time estimates are in general agreement with prior dated tree estimates but with the current study finding slightly older estimates (17, 43).

**Figure 1.**
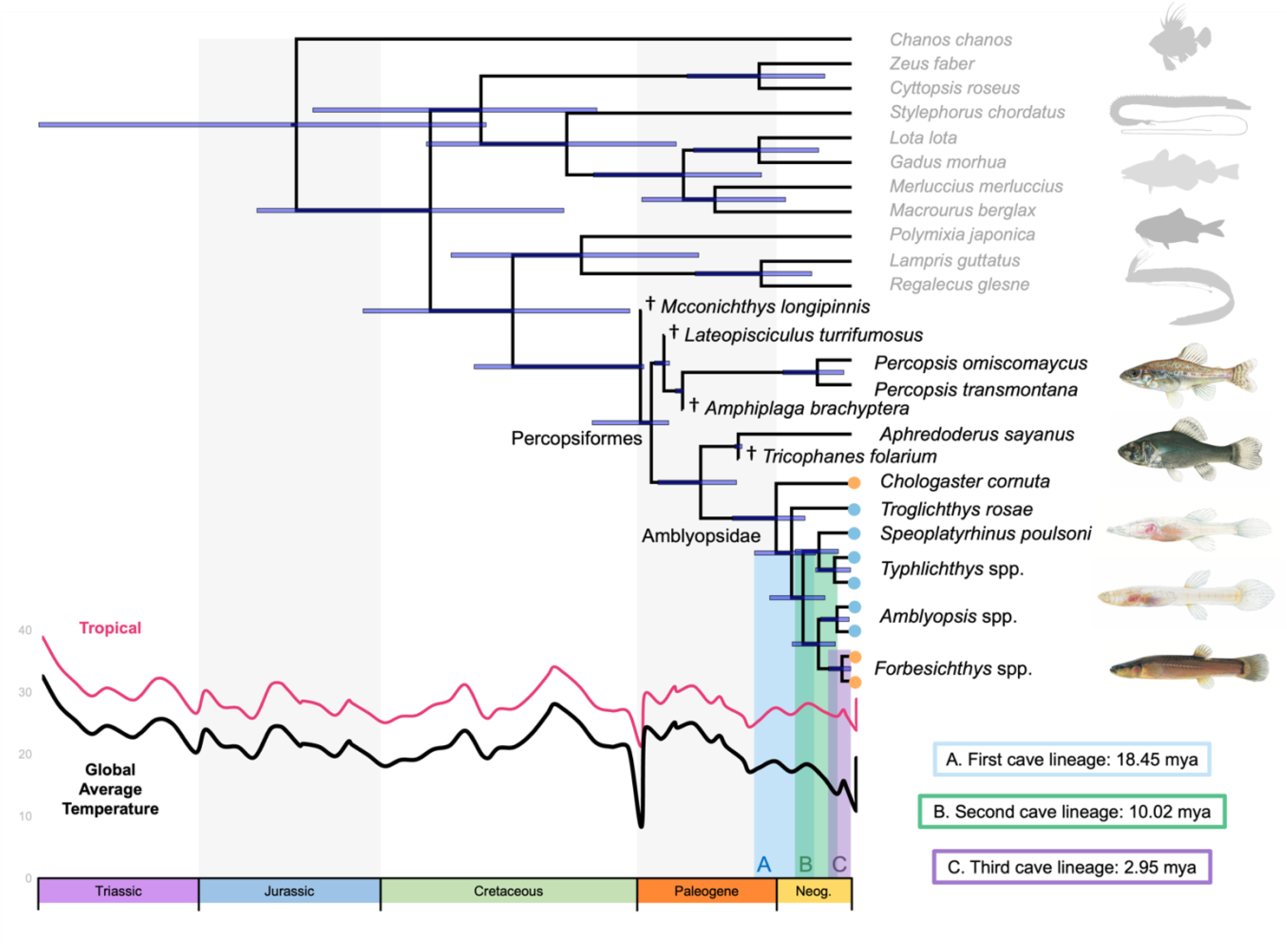
Time-calibrated phylogeny of the Percopsiformes with four fossil taxa, all extant genera, and most species. Three independent cave invasion events are indicated with vertical bars below the phylogeny: first invasion 11–29.8 Mya (18.5 median, ice blue bar), 4.1–17.3 Mya (10.0 median, green bar), and 0.3 to 7.1 Mya (3.0 median, purple bar). Tropical and Global Average temperature curves used in evolutionary model testing are also included below the phylogeny with degrees Celsius on the x-axis. Horizontal boxes on each node indicate the 95% HPD. Cave (blue) and non-cave (orange) fishes of the Amblyopsidae are denoted with circles at the phylogeny tips. Fossil taxa are distinguished by †. For upper and lower 95% HPD ages, see Supplementary Material 2 Fig. S1. Illustrations from top to bottom: *Percopsis transmontana*, *Aphredoderus sayanus*, *Speoplatyrhinus poulsoni*, *Typhlichthys subterraneus*, and *Forbesichthys agassizii*. Illustration copyrights by Joseph Tomelleri.

The fossil record provides context for our phylogenetic findings. Amblyopsidae fossils are scarce, with some vertebrae tentatively assigned to the family from the Late Eocene (37.8– 33.9 Mya) – Early Oligocene (33.9–28.1 Mya) Cypress Hills formation of Saskatchewan and Alberta, Canada (45). However, other percopsiforms have a more extensive fossil record. *Tricophanes* from the Oligocene, likely sister to *Aphredoderus*, shares caudal fin modifications (46) as well as strongly serrate infraorbital bones (JWA pers. obs.). Percopsidae has an extensive fossil record with several described fossil genera (47), with the oldest articulated percopsiform, *Lindoeichthys*, dating from the Maastrichian. Additional dentaries from the Campanian (late Cretaceous) indicate that there were already multiple percopsiform species present by ∼83.6– 72.1 Mya (47).

### Paleoclimatic influence on cave colonization

Using a macroevolutionary model fitting approach, we found that transitions from surface to cave habitats in percopsiform fishes are best explained by a paleotemperature-driven model of evolution. We explored three potential modes of cave colonization: paleotemperature-dependent models (climate-relict hypothesis) (Melendez-Vazquez et al., under review), early burst adaptive radiation hypothesis, and Brownian motion (stochastic colonization hypothesis). The climate model received the best-fit to our data in all analyses (Table 1) with the Global Average Temperature Scale yielding an AICw = 0.8710 and the Tropical Scale an AICw = 0.8419.

**Table 1.** Support for evolutionary model testing using two climate-curves (48) run for 1000 iterations. Bolded values indicate the model with the best fit for each climate-curve tested.

| Facultative Cave-dweller as: | Scale | Climate Model Fit AICw | Brownian Motion Model Fit AICw | Early Burst Model Fit AICw | Climate Model Fit AIC | Brownian Motion Model Fit AIC | Early Burst Model Fit AIC | Beta |
| --- | --- | --- | --- | --- | --- | --- | --- | --- |
| Non-cave | GAT | **0.8710** | 0.0752 | 0.0536 | **51.3166** | 56.2134 | 56.8900 | 18.0282 |
| Non -cave | Tropical | **0.8419** | 0.0922 | 0.0657 | **51.7910** | 56.2134 | 56.8900 | 15.9338 |
| Cave | GAT | **0.9359** | 0.0233 | 0.0407 | **34.3344** | 41.7160 | 40.6027 | 21.4941 |
| Cave | Tropical | **0.8684** | 0.0479 | 0.0836 | **35.9215** | 41.7160 | 40.6027 | 20.0044 |

The timing of each cave colonization event, as estimated from our phylogenetic analysis, does not correspond to major extinction events such as the Cretaceous-Paleogene Extinction Event (K–Pg, 66 Mya) or the Paleocene-Eocene Thermal Maximum (PETM, 56 Mya), both of which had extensive effects on global biodiversity (49–51). Instead, our results indicate that climatic events following the Early-Middle Eocene cooling (47 Mya) may have been more influential. According to benthic δ^18^O values, two notable climatic transitions occurred between 34 Mya to present: a slight warming until the Middle Miocene Climate Transition at around 14 Mya and the last major climatic transition that led to the Pleistocene and glaciation of the Northern Hemisphere at 2.8 Mya (52). From 34 Mya to the present day, more positive benthic δ^18^O values are associated with colder climate conditions, a progressive decrease in CO_2_, lower global mean sea-level, and lower carbonate concentration depth (52).

These findings support the climate-relict hypothesis, which posits that cave colonization may have been a response to changing surface conditions. As surface habitats became less favorable due to climate shifts, pre-adapted populations may have found refuge in more stable cave environments. This mechanism has been suggested as a mode of cave colonization leading to troglobiotic organisms, where climate-relict populations in caves remain and establish following a climate transition that decimates the necessarily pre-adapted surface populations (21, 22).

However, our results also suggest that climate alone may not fully explain the pattern of cave colonization. The potential influence of biotic factors, such as competition or predation, should also be considered. Interestingly, the timing of the leuciscid clade migration to North America (30–37 Mya) (45, 53) is close to our uppermost cave colonization estimate for percopsiforms (29.8 Mya). This temporal proximity raises the possibility that competitive pressures from newly arrived fish groups may have contributed to cave colonization by pre-adapted percopsiform ancestors.

While our model testing strongly supports paleoclimatic shifts as a primary driver of cave colonization in North American cavefishes, it is important to note that alternative hypotheses have been supported in other systems. For instance, support for stochastic cave colonization events has been found in Hawaiian planthoppers (28) and the Mexican Blind Cave Tetra (27, 54), as well as biologically driven adaptive shifts in Brazilian planthoppers (55). These examples underscore the complexity of cave colonization processes and the potential for different mechanisms to dominate in different systems.

### Comparative genomics of troglomorphic traits

We examined three hypotheses, which either included (Scheme 1), excluded (Scheme 2), or specifically referenced the facultative cave-dwelling fishes (Scheme 3). In our comparative genomic analyses, we identified selective regime shifts in 226 loci associated with troglomorphy, observing both relaxed and intensified selection. Interestingly, we found different genes related to the same trait (e.g., pigmentation genes *bnc2* and *mc1r*) to be under both relaxed and intensified selection. Of the genes analyzed, only two (*gnb1a* and *mitfa*) shifted the direction of selection depending on the specific hypotheses tested.

Out of the 226 loci analyzed for signatures of intensified or relaxed selection, 50 were under either relaxed or intensified selection in Scheme 1, 58 in Scheme 2, and 51 in Scheme 3. Consistently, we found more loci under relaxed selection than intensified. Our first hypothesis (Scheme 1, Fig. 2) included facultative cave-dwelling species as reference branches, cave-obligate species as test branches, and all other species as background branches. There were 14 genes undergoing intensified selection (GUIS) found from the test branches with 25 GUIS from the reference branches, with 29% (eight loci) of GUIS shared between the test and reference branches. We found several more genes undergoing relaxed selection (GURS) in comparison to GUIS, with 36 GURS from the test branches and 25 GURS from the reference branches. The test and reference branches shared 45% (19 genes) of GURS.

**Figure 2.**
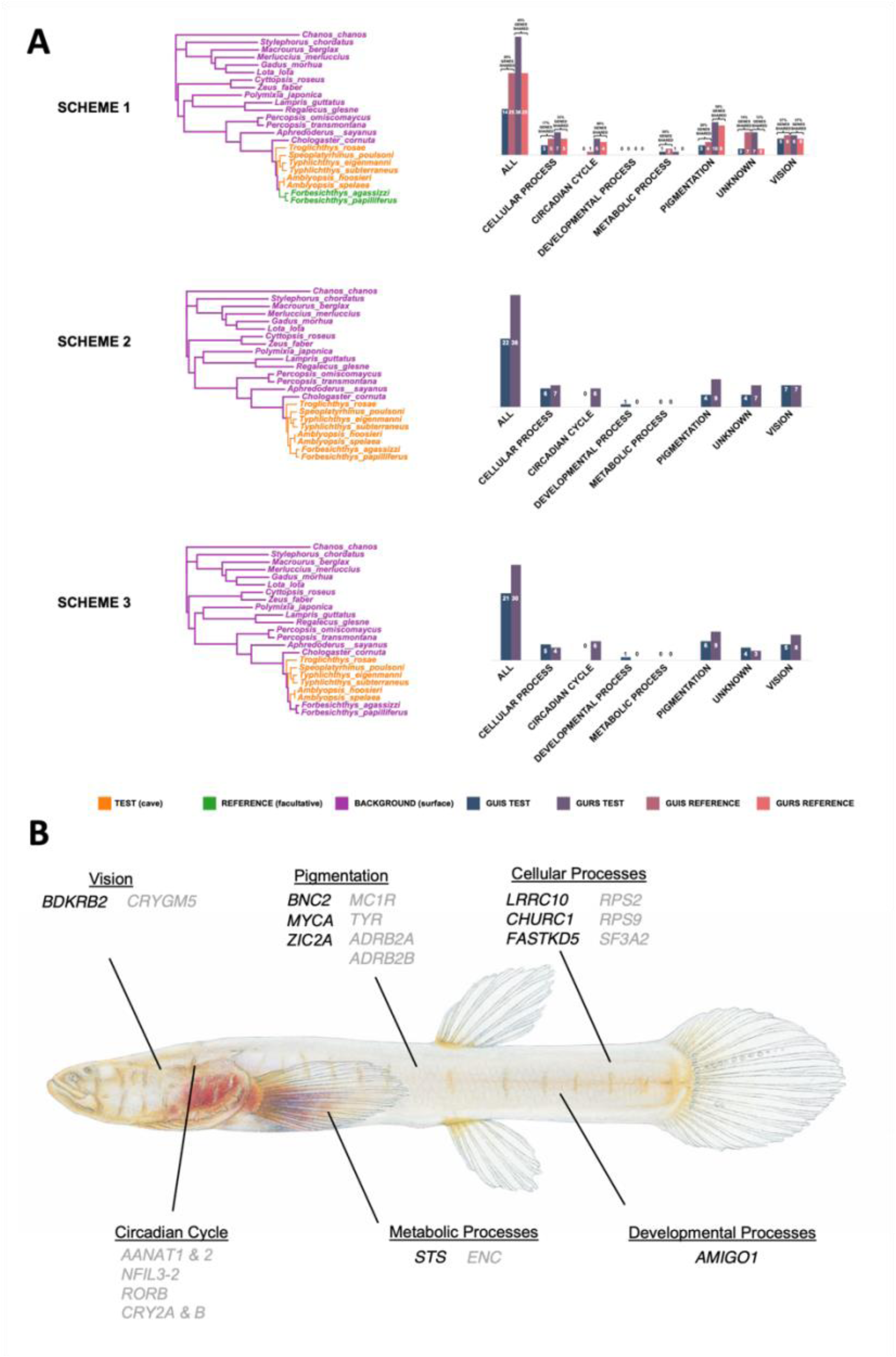
Results of HyPhy RELAX analyses indicating genes under intensified and relaxed selection in all three cave-obligate designation schemes (A). An arbitrary selection of genes with the same selection intensity across schemes, showing intensified (black) and relaxed (grey) selection, labeled with PANTHER estimated functions (B); Full list of candidate genes can be found in Supplementary Material S3. GUIS stands for genes under intensified selection; GURS stands for genes under relaxed selection. Illustration copyright by Joseph Tomelleri, *Typhlichthys subterraneus*.

Our second hypothesis included the two species of facultative cave-dwelling fishes within the test group with all other species as background branches, while our third hypothesis had only cave-obligate fishes as test branches. The greatest number of GUIS for Scheme 2 were related to vision (seven loci) and for Scheme 3 were pigmentation (six loci). The greatest number of GURS for both Schemes 2 and 3 were related to pigmentation (nine loci). For GUIS, we found 14 genes shared between Schemes 2 and 3 (Supplementary Material S3–S6). When comparing Schemes 2 and 3, nine GUIS were unique to Scheme 2 and five GUIS were unique to Scheme 3. We found 24 GURS shared between Schemes 2 and 3 (Supplementary Material S3– S6). We found 11 unique GURS for Scheme 2 and 7 unique GURS for Scheme 3 when comparing the two hypotheses.

### Selective regime shifts in cave-dwelling fishes

We found evidence for both relaxed and intensified selection in our troglomorphic candidate loci across multiple schemes with cave, non-cave, and reference (facultative cave-dwelling) species (Fig. 2A). We found signals of relaxed selection in some pigmentation (e.g., *mc1r* and *tyr*) and vision-related (e.g., *crygm5*) loci for cave amblyopsid fishes across our schemes (Fig. 2B, Supplementary Material S3). Relaxation of selection can occur due to the absence of a functional constraint or a decrease in the efficiency of selection (56, 57); neutral mutations may then accumulate and render a gene ineffectual (58, 59). The lack of light in caves removes most functional constraints of pigment (i.e., colouration for breeding, UV protection, and crypsis) and vision.

The *tyr* locus is interesting as it has been shown to lead to albinism in zebrafish, mice, rabbits, and ducks when down regulated (60–63); and changes to *tyr* (as well as *tyro1* and *mpv17*) led to the loss of iridiophores in the Chinese cavefish *Sinocyclocheilus* (64). Li et al. (65) did not find any differences in *tyr* in surface vs. cave *Sinocyclocheilus* but did find down regulation in a crucial gene that leads to activation of the *tyr* promoter. Loss-of-function mutations in *mc1r* in zebrafish leads to no changes in regular pigmentation, but a loss of countershading, which is a critical part of colouration that is unnecessary in the cave environment (66).

Our analyses suggest that for amblyopsids, the removal of these functional constraints may result in relaxation of selection for some pigmentation and vision-related loci. Though macroevolutionary models have not been applied directly to other cavefishes, additional work has shown mutations in several troglomorphic loci (as well as expression changes and other genomic structural variations) that may also bear weight towards neutral mutation as a mechanism of troglomorphy (64).

We also detected signals of intensification of selection for other vision (e.g., *bdkrb2* and *opn3*) and pigment (e.g., *bnc2* and *zic2a*) loci across our branch-site model schemes (Fig. 2B, Supplementary Material S3). Moran et al. (67) suggests that the loss of eyes in cave animals is itself an adaptive response (i.e., directional selection for lack of eyes) due to the high metabolic cost of eyes and optic lobes. The metabolic cost to an organism creating and maintaining an eye and optic lobes can be staggering (67). There may also be a constructive function for the loss of eyes, as their loss leads to a greater area for mechanosensory hair cell units (neuromasts) to develop (68). As in Mexican Blind Cave Tetras, lines of neuromasts occur where the eyes would have been in cave obligate amblyopsids (39, 69). Thus, the intensified selection on vision and pigment related genes may be due to an increase in selection (i.e., directional) against those tissues and functions and for increased space for sensory hair cells.

The discovery of both relaxed and intensified selection signals in loci relating to the same function or process indicates the complex nature and interplay of selective pressures in differing selective regimes or could point to additional roles of these loci in other aspects of the life cycle of these fishes. This pattern suggests that while some aspects of these traits may be under reduced selective pressure in the cave environment, others may be co-opted for new functions or modified to suit the unique demands of cave life.

Overall, our study sheds light on the evolutionary history and adaptive processes of cave colonization in North American cavefishes, revealing key periods of colonization driven by paleoclimatic shifts in the Miocene and Pliocene. These findings not only deepen our understanding of adaptation to extreme environments but also offer insights relevant to broader fields such as conservation and medical research. Our analysis of the genomic basis of cave adaptation in amblyopsid fishes has important implications for evolutionary biology, demonstrating that the evolution of cave-dwelling traits is not simply a matter of trait loss, but rather a nuanced process of trait modification and potential co-option for new functions.

The observed shifts in selective regimes across different loci highlight the mosaic nature of genomic evolution during habitat transitions. While some pigmentation and vision-related genes (e.g., *mc1r*, *tyr*, *crygm5*) show signals of relaxed selection, others (e.g., *bdkrb2*, *opn3*, *bnc2*, *zic2a*) exhibit intensified selection, underscoring the complex interplay of selective pressures acting differentially across the genome. Our results also contribute to the ongoing debate about the relative importance of neutral versus adaptive processes in the evolution of cave organisms. The prevalence of relaxed selection in many troglomorphic loci supports the idea that neutral processes play a significant role in cave evolution, while the detection of intensified selection in other loci suggests that adaptive processes are also at work, potentially fine-tuning or repurposing existing traits for the cave environment.

## Materials and Methods

### Taxonomic sampling

Muscle tissue or fin clip samples were obtained for all percopsiform genera (6 genera) and all amblyopsid species (13 species in total), including two of five newly revised *Aphredoderus* species (*Ap. sayanus* and *Ap. gibbosus*) (36). For the phylogenomic analyses, we sequenced multiple individuals per species (n= 2–4) for nearly all (11) species (42 specimens). However, due to their highly protected status, we sequenced only single individuals for the Alabama (*Speoplatyrhinus poulsoni*) and Ozark (*Troglichthys rosae*) Cavefishes. Our sampling spans the range of troglomorphy in the family Amblyopsidae: surface-dwelling (the Swampfish, *Chologaster cornuta*), facultative cave-dwelling (or surface-cave, the Spring Cavefishes, *Forbesichthys* spp.), and cave-obligate fishes (the Northern Cavefishes, *Amblyopsis* spp., the Southern Cavefish, *Typhlichthys subterraneus*, and the aforementioned Alabama and Ozark Cavefishes). See details in supplementary Table 1.

### Genomic data collection

For eleven percopsiform species (n = 31 individuals), DNA was extracted from muscle or fin clip using DNeasy Blood and Tissue Kits (Qiagen). To expand our taxonomic sampling, we included previously extracted DNA samples from Niemiller et al. (43) and Hart et al. (38) for two additional species and additional replicates (11 specimens). Library preparation for whole-genome short-read sequencing was outsourced to ArborBiosciences using the Illumina Nextera DNA sample preparation kit. Libraries were sequenced on NovaSeq S4 (generating up to 100 million reads pair-ended per sample). For quality control purposes and to facilitate genome assembly, we sequenced multiple individuals per species (n= 2–4) to identify potential instances of contamination and facilitate genome assemblies. To account for the phylogenetic uncertainty regarding the sister-group relationship with Percopsiformes, we expanded our dataset using published data for six fish orders from National Center for Biotechnology Information (NCBI) (Supplementary Material S1). Raw sequence reads were deposited on NCBI under BioProject PRJNA737769, PRJNA737771, PRJN737782 and some under review (accession numbers will be available in the Supplementary Material).

### Whole-genome short-read assembly and exon marker extraction

Raw Illumina reads were assessed for quality using FastQC (v0.11.5) (70) and trimmed using Trimmomatic (v0.38) (71) to remove low-quality bases and adapters. The resulting high-quality reads were then assembled using SPAdes (v3.13.1) (72). To mine single-copy exon markers from the percopsiform whole-genome short-read data, we utilised the FishLifeExonHarvesting pipeline (73), with the Backbone 2 probe set containing 1,105 markers. Hidden Markov Model (HMM) profiles were generated for each exon and searched against the genome assemblies using HMMER (v.3.2.1) (48) with a bit score cut-off of 100. Identified hits were concatenated and converted to FASTA format using a custom Python script. Reading frame correction was performed with CD-HIT (74) and exon filtering was carried out using Exonerate (75) to ensure proper exon boundaries. Finally, all harvested exons were aligned with MACSE (v2.03) (76), which considers reading frames, using a custom bash script.

### Phylogenomic estimation

All gene alignments were concatenated into a single super-matrix using the AMAS package (77). To quickly identify potential instances of contamination or misidentification, we estimated an initial tree using FastTree (v2.0) (78). After this initial quality control step, we refined our matrix to include only one representative per species to minimise missing data. We then estimated a maximum likelihood tree in IQ-TREE (v2.0) (79), employing mixture models to account for heterogeneity in evolutionary rates across sites. We assessed branch support using the nonparametric ultrafast bootstrap (UFBoot2) algorithm. To account for incomplete lineage sorting (ILS), we performed a multiple-species coalescent species trees analysis. Gene trees were estimated using the aforementioned IQ-TREE parameters, which served as input data for ASTRAL-III (80). Prior to their use in ASTRAL-III, poorly supported relationships in individual gene trees (bootstrap support ≤ 33) were collapsed to minimise estimation errors (80).

### Integration of fossil and extant species

To calibrate our phylogeny and incorporate fossil lineages, we used fossil age estimates and original descriptions for extinct Percopsiformes from the Paleobiology Database (https://paleobiodb.org). Four fossil taxa were selected based on available morphological data: *Mcconichthys longipinnis*† Grande 1988*, Lateopisciculus turrifumosus*† Murray and Wilson 1996*, Amphiplaga brachyptera*† Cope 1877, and *Tricophanes folarium*† Cope 1872. We compiled literature to score and place these fossils into the morphological matrix from Armbruster et al. (42), including them as terminal taxa (Supplementary Material S2) (46, 81–87). Our final morphological dataset consisted of 66 skeletal characters, encompassing four fossils and 13 extant percopsiform species (nine amblyopsid, two aphredoderid, and two percopsid species). This combined dataset of fossil and extant taxa allowed us to jointly estimate the phylogenetic placement of fossils and divergence times.

### Divergence time estimation

We generated a time calibrated phylogenomic hypothesis using the Fossilised Birth Death (FBD) model in MrBayes (v.3.2.7a) (88). For this analysis, we assembled a subset of exon loci based on the mean coefficient of rate variation (a measure of clock-likeliness), totaling 241 clock-like loci via PAUP (v.4.0a) (89). To minimise missing data, we pruned our extended phylogeny to one individual per species based on the completeness of sequence data. Our genomic data was combined with our morphological dataset containing both fossil and extant taxa. We partitioned our molecular dataset by codon position using the General Time Reversible model with gamma-distributed rate variation across sites (GTRGAMMA). The morphological data was analyzed using the Mk model for discrete morphological characters. We employed a relaxed clock model with a lognormal distribution clock rate prior and an independent gamma rate (IGR) model to account for rate variation among lineages. Convergence of the Markov chain Monte Carlo (MCMC) analyses was determined by an effective sample size (ESS) value close to or above 200. Tree sampling occurred every 10,000 generations. A list of the fossil ages is provided in Supplementary Material S2.

### Paleotemperature data and evolutionary model fitting

To evaluate the association between cave colonization and ancient climate changes, we applied a recently proposed threshold model (Melendez-Vazquez et al., under review) to examine the relationship between cave and non-cave lineages (as a discrete trait) and historical climatic variations, represented by a time-dependent curve. This model builds upon a likelihood approximation achieved through numerical integration and quadratures, following the general algorithm outlined by Hiscott et al. (90). By adapting this climatic threshold model, we were able to employ a paleoclimatic scale to ascertain if ancient cold or warm periods have instigated transitions to cave environments. We fitted previously published paleoclimatic scales (48) to our model and assigned character states (cave or non-cave) for each taxon based on their habitats. Facultative cave-dwelling fishes were categorised as both non-cave or cave-dwellers, and separate analyses were conducted to account for trait assignation uncertainty. We conducted a model selection analysis comparing the fit of our modified climatic threshold model (CLIM) to other models, including Brownian Motion (BM) and Early Burst (EB). Our CLIM model allows the rate of evolution to vary at time points corresponding to paleotemperature changes. These analyses were conducted using 1,000 integrations (n = 1,000) with a climatic spline interpolation function of 500 degrees of freedom (df = 500). We use the global average paleo-temperature curve (48), along with additional paleo-climatic curves based on tropical and global temperatures, to enhance the robustness of our evolutionary analyses. All evolutionary model fitting analyses were conducted in R (v4.2.1).

### Comparative genomics and in silico capture of candidate genes

We used a hidden Markov model (HMM) approach in HMMER v3.2.1 (91) to identify and extract 458 candidate genes from the assembled genomes, based on previous studies related to vision, pigmentation, and circadian cycles in cavefishes and subterranean mammals (92–95). Of these, 398 were successfully mined, and we incorporated an additional 89 single-copy genes identified using BUSCO (Actinopterygii OrthoDB) and OrthoFinder (96, 97), totaling 487 genes. Sequence alignments were performed using MACSE (76) and visually inspected with Geneious Prime® v2022.1. We then applied filters to retain genes present in at least 70% of species, with at least 70% sequence similarity and length greater than 200 bp, resulting in 226 genes (73 from OrthoFinder, 153 from previous studies) for downstream analyses. Functions and pathways of BUSCO genes were determined using PANTHER (v2.2) (98), associating focal genes with processes including cellular functions, circadian cycles, development, metabolism, pigmentation, vision, and genes without ascribed functions.

### Assessment of relaxed and intensified selection

We employed the RELAX framework (57) implemented in the package HyPhy (v2.5.58) (99) to investigate signatures of intensified or relaxed selection in loci associated with vision, pigmentation, circadian cycles, hematopoiesis, and other functions. We compared foreground branches (test and reference, Fig. 2) of cavefishes to background (non-cavefish) branches, based on estimates of the selection strength parameter *K* (using a *p* value = 0.05). A value of *K* < 1 indicates that test branch was under significant relaxed selection, whereas a value of *K* > 1 suggests intensified selection on the test branches. We applied a correction to the p-values using the *p.adjust* tool and the Benjamini-Hochberg procedure using the R package *stats* (stats v3.6.2) with a cut-off of 0.05. We tested three alternative schemes: (1) obligate cavefish species (*Tr. rosae, S. poulsoni, Ty. eigenmanni, Ty. subterraneus, A. hoosieri,* and *A. spelaea*) as test branches, facultative cavefish species (*F. agassizii* and *F. papilliferus*) as reference branches, and remaining species as background; (2) both obligate and facultative cavefish species as test branches, with remaining species as background; and (3) obligate cavefish species as test branches, with facultative cavefish and remaining species as background (Fig. 2). These comparative analyses across different evolutionary stages of cave adaptation allowed us to elucidate the molecular underpinnings of troglomorphic traits and to assess the relative contributions of relaxed and intensified selection in shaping cave-adapted genomes.

## Acknowledgments

We would like to acknowledge the Louisiana State University Museum of Natural Science Tissue Collection, the North Carolina State Museum, the Ohio State University Museum Tissue Collection, the University of Tennessee Tissue Collection, and the Yale Fish Tissue Collection for their museum tissue specimen loans. We would also like to thank our funding resources: The National Science Foundation Postdoctoral Research Fellowship in Biology to P.B. Hart and the University of Oklahoma and Sam Noble Oklahoma Museum of Natural History to D. Arcila.

## CRediT Author Contributions

<u>Conceptualization</u>: P.B. Hart, D. Arcila.

<u>Data Curation</u>: P.B. Hart, D. Arcila, M. Rincon-Sandoval, F. Melendez-Vazquez, O. Bierstein, B. Gough, E. M. Troyer

<u>Formal Analysis</u>: M. Rincon-Sandoval, D. Arcila, F. Melendez-Vazquez, P.B. Hart, E. M. Troyer

<u>Funding Acquisition</u>: D. Arcila, P.B. Hart

<u>Investigation</u>: P.B. Hart, D. Arcila, M. Rincon-Sandoval

<u>Methodology</u>: M. Rincon-Sandoval, D. Arcila, F. Melendez-Vazquez, E. Troyer, P.B. Hart, E. M. Troyer

<u>Resources</u>: D. Arcila, P.B. Hart, M.L. Niemiller, J.W. Armbruster

<u>Software</u>: M. Rincon-Sandoval, D. Arcila, F. Melendez-Vazquez

<u>Supervision</u>: P.B. Hart, D. Arcila

<u>Validation</u>: M. Rincon-Sandoval, D. Arcila, F. Melendez-Vazquez, P.B. Hart

<u>Visualization</u>: P.B. Hart, M. Rincon-Sandoval

<u>Writing – Original Draft</u>: P.B. Hart, D. Arcila, M. Rincon-Sandoval

<u>Writing – Review & Editing</u>: D. Arcila, P.B. Hart, M.L. Niemiller, J.W. Armbruster, R. Betancur-R, E. Troyer, M. Rincon-Sandoval, F. Melendez-Vazquez, O. Bierstein, B. Gough.

