## Supplementary Material 1 and 2 for "Ancient climate changes and relaxed selection shape cave colonization in North American cavefishes"

**Supplementary Material S1 for "Ancient climate changes drive mode and tempo of cave colonization in North American cavefishes"**

### Supplementary Table

**Table S1** Tissue specimens used for whole shotgun genome resequencing. Museum codes are: LSUMNS: Louisiana State University Museum of Natural Science, NCSM: North Carolina State Museum, UTTC: University of Tennessee Tissue Collection, YFTC: Yale Fish Tissue Collection.

| Species | BioProject | BioSample | Sample No. | Type | State | County | Site |
|---|---|---|---|---|---|---|---|
| *Amblyopsis hoosieri* | PRJNA737769 | SAMN20209477 | MLN 0246 | Tissue | IN | Lawrence | Donaldson Cave |
| *Amblyopsis hoosieri* | PRJNA737769 | SAMN20209476 | MLN 0243 | Tissue | IN | Lawrence | Blue Springs Caverns |
| *Amblyopsis hoosieri* | PRJNA737769 | SAMN20209475 | MLN 0242 | Tissue | IN | Lawrence | Donaldson Cave |
| *Amblyopsis spelaea* | PRJNA737771 | SAMN20209478 | YFTC 23868 | Extract | KY | Breckinridge | Webster Cave |
| *Amblyopsis spelaea* | PRJNA737771 | SAMN20209479 | YFTC 23869 | Extract | KY | Breckinridge | Webster Cave |
| *Amblyopsis spelaea* | PRJNA737771 | SAMN20209480 | YFTC 23875 | Extract | KY | Breckinridge | Webster Cave |
| *Aphredoderus sayanus* | Under review | Under review | LSUMNS 3030/LSUMNZ 15287 | Extract | LA | Bienville Parish | 5 mi W Lucky |
| *Aphredoderus sayanus* | Under review | Under review | TJN 244 | Extract | NC | Nash | Swift Creek |
| *Aphredoderus sayanus* | Under review | Under review | OSUM 113433-1 | Tissue | OH | Allen | Auglaize River |
| *Aphredoderus sayanus* | Under review | Under review | OSUM 113433-2 | Tissue | OH | Allen | Auglaize River |
| *Chologaster cornuta* | Under review | Under review | YFTC 23849 | Extract | NC | Bladen | Colly Creek at NC 53 |
| *Chologaster cornuta* | Under review | Under review | NCSM 47776 | Extract | NC | Cumberland | Unnamed tributary, north of Shaw Road [SR 1437] at Bonnie Doone Lake, [ca. 5.0 air miles NW center Fayetteville] |
| *Chologaster cornuta* | Under review | Under review | NCSM 83831 | Extract | NC | Cumberland | [Indian Creek], outflow stream from Jessups Pond, on West side of [SR]53, ca. 1 kilometer N of Bladen Country line, [ca. 28.3 kilometers NNW center Elizabethtown] |
| *Chologaster cornuta* | Under review | Under review | DAN 286 | Extract | NC | | |

| | | | | | | | |
|---|---|---|---|---|---|---|---|
| *Forbesichthys agassizzi* | Under review | Under review | MLN RP2 | Extract | TN | Coffee | Rigsby Pond |
| *Forbesichthys agassizzi* | Under review | Under review | UTTC 242 | Extract | TN | DeKalb | Blue Springs |
| *Forbesichthys agassizzi* | Under review | Under review | UTTC 243 | Extract | TN | DeKalb | Blue Springs |
| *Forbesichthys agassizzi* | Under review | Under review | DAN 282 | Extract | TN | Coffee | Fultz Pond |
| *Forbesichthys agassizzi* | Under review | Under review | UTTC 649 | Extract | TN | DeKalb | Blue Springs |
| *Forbesichthys papilliferus* | Under review | Under review | 23877 | Extract | KY | Todd | Spring-fed ditch N of crossing with Morton Rd |
| *Forbesichthys papilliferus* | Under review | Under review | 23878 | Extract | KY | Todd | Spring-fed ditch N of crossing with Morton Rd |
| *Forbesichthys papilliferus* | Under review | Under review | UTTC 102 | Extract | IL | Union | Cave Spring Cave |
| *Forbesichthys papilliferus* | Under review | Under review | FPAP-CLK1 | Tissue | TN | Montgomery | Clarksville Lake Cave |
| *Forbesichthys papilliferus* | Under review | Under review | TJN 100 | Extract | IL | Union | Cave Spring Cave |
| *Percopsis omiscomaycus* | Under review | Under review | TJN 237 | Extract | IL | Putnam | Illinois River at Hennepin |
| *Percopsis omiscomaycus* | Under review | Under review | OSUM 113977-1 | Tissue | OH | Franklin | Big Walnut Creek |
| *Percopsis omiscomaycus* | Under review | Under review | OSUM 113977-2 | Tissue | OH | Franklin | Big Walnut Creek |
| *Percopsis transmontana* | Under review | Under review | T1890 | Tissue | WA | Skamania | |
| *Percopsis transmontana* | Under review | Under review | T1894 | Tissue | WA | Skamania | |
| *Percopsis transmontana* | Under review | Under review | T1895 | Tissue | WA | Skamania | |
| *Speoplatyrhinus poulsoni* | Under review | Under review | PBH-Speo01-2019 | Extract | AL | Lauderdale | Key Cave |
| *Troglichthys rosae* | PRJNA737782 | SAMN20209507 | L1 | Extract | AR | Benton | Logan Cave |
| *Troglichthys rosae* | PRJNA737782 | SAMN20209506 | C5 | Extract | AR | Benton | Cave Springs Cave |
| *Troglichthys rosae* | PRJNA737782 | SAMN20209505 | C3 | Extract | AR | Benton | Cave Springs Cave |
| *Typhlichthys eigenmanni* | Under review | Under review | MLN 0414 | Tissue | MO | Camden | Carroll Cave |
| *Typhlichthys eigenmanni* | Under review | Under review | MLN 0404 | Tissue | MO | Shannon | Flying W Cave |
| *Typhlichthys eigenmanni* | Under review | Under review | MLN 0429 | Tissue | MO | Oregon | Falling Spring Cave |
| *Typhlichthys eigenmanni* | Under review | Under review | EN1 | Extract | AR | Stone | Ennis Cave |
| *Typhlichthys subterraneus* | Under review | Under review | MLN 0272 | Tissue | TN | Putnam | Stamps Cave |
| *Typhlichthys subterraneus* | Under review | Under review | MLN 0296 | Tissue | TN | Decatur | Baugus Cave |
| *Typhlichthys subterraneus* | Under review | Under review | MLN 0051 | Tissue | TN | Coffee | Blowing Springs Cave |
| *Typhlichthys subterraneus* | Under review | Under review | MLN 18-035.3 | Specimen | AL | Jackson | Fern Cave |

**Supplementary Material S2 for "Ancient climate changes drive mode and tempo of cave colonization in North American cavefishes"**

**Fossil Calibration Age Priors & Integration of Fossil and Extant Species**

†*Mcconichthys longipinnis* Grande 1988, Mcconichthyidae, Percopsiformes. Calibration ages from Alfaro et al. (2018), min age: 63.1, max age: 93.51. Three phylogenetic placements for †*Mcconichthys longipinnis* are found in the literature, with the fossil either being a deeply nested within Paracanthopterygii (Grande 1988), a stem percopsiform (Murray and Wilson 1999) or an aphredoderid (Borden et al. 2013). †*Mcconichthys longipinnis* was conservatively placed as a stem percopsiform based on the characters of foreshortened centra and all hypurals separate from ural centrum 2 (Murray and Wilson 1999, Grande et al. 2013) and was scored on characters 52, 54, 55, and 56 (Murray and Wilson 1999, Borden et al. 2013, Grande et al. 2013, Armbruster et al. 2016).

†*Lateopisciculus turrifumosus* Murray and Wilson 1996, Percopsidae, Percopsiformes. Calibration ages from Murray and Wilson (1996), min age: 56, max age: 66. We placed †*Lateopisciculus turrifumosus* within Percopsidae based on the presence of a dorsal process on the maxilla and the presence of six autogenous hypurals, the latter of which unites fossil percopsids (Murray and Wilson 1996, Murray and Wilson 1999, Borden et al. 2013). We scored the fossil on characters 52–56 (Murray and Wilson 1996, Murray and Wilson 1999. Borden et al. 2013, Armbruster et al. 2016).

†*Amphiplaga brachyptera* Cope 1877, Percopsidae, Percopsiformes. Calibration ages from Newbrey et al. (2013) and Grande et al. (2013), min age: 50.3, max age: 55.8. †*Amphiplaga*

*brachyptera* has been consistently placed within Percopsidae with the character of six autogenous hypurals (uniting all fossil percopsids) and two epurals (Murray and Wilson 1999, Borden et al. 2013, Grande et al. 2013, Newbrey et al. 2013). We also placed the fossil in Percopsidae and scored characters 52–56 (Murray and Wilson 1999, Borden et al. 2013, Grande et al. 2013, Newbrey et al. 2013, Armbruster et al. 2016).

†*Tricophanes folarium*, Aphredoderidae, Percopsiformes. Calibration ages from Grande et al. (2013) and Near et al. (2012), min age: 33.7, max age: 36. †*Tricophanes folarium* and Aphredoderus are united by having fused hypurals 3–5 to each other and ural centrum 2 with a distinct hypural 6 (Grande et al. 2013, Armbruster et al. 2016).

### Supplementary Figures

**Figure S1.** Time-calibrated total-evidence phylogenetic hypothesis with four fossil and all extant genera of the Percopsiformes fishes. Horizontal node bars indicate the 95% HPD. The upper and lower age limits of the 95% HPD are included directly above the corresponding node bar.

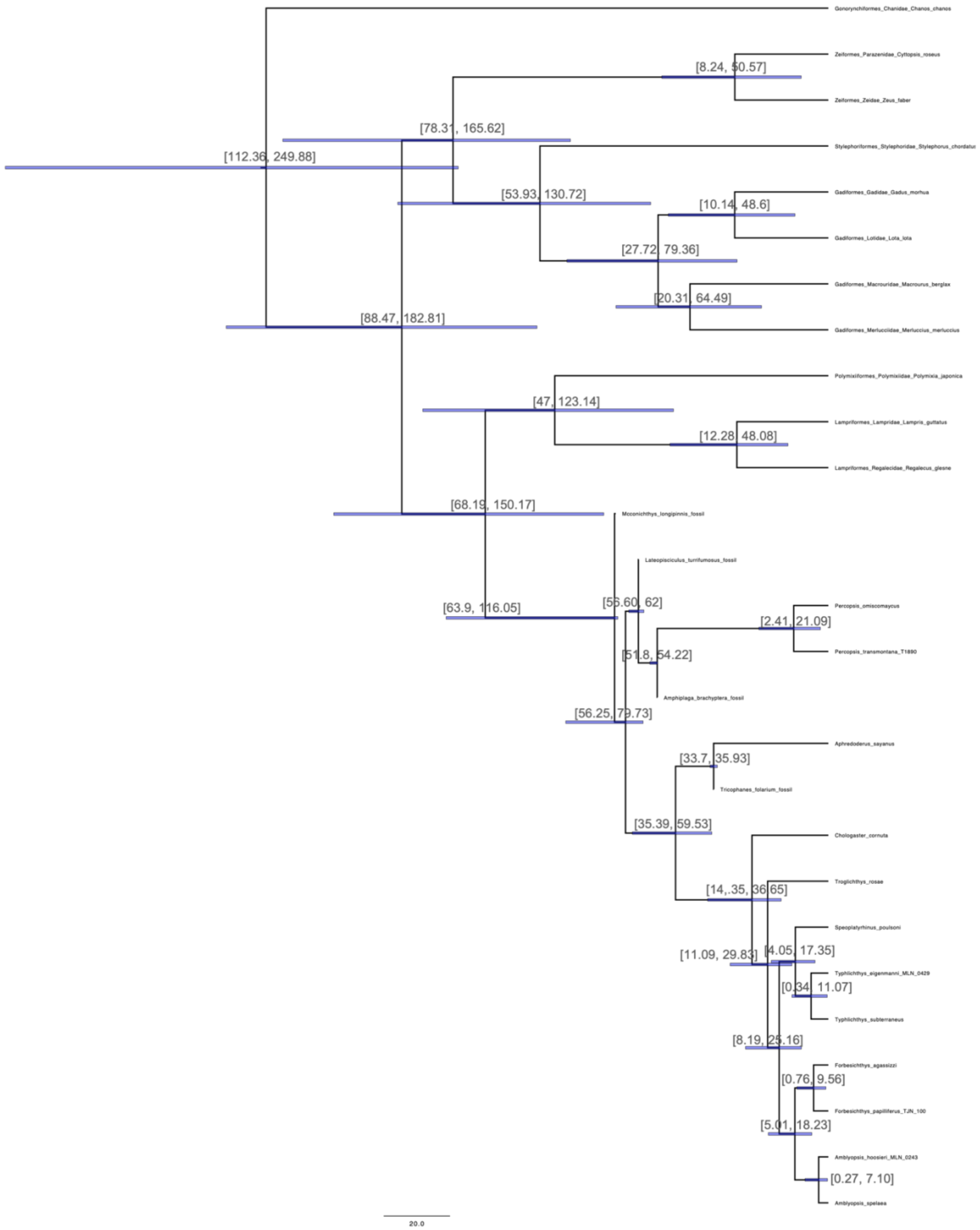
