## Supplementary Material 6 for "Ancient climate changes and relaxed selection shape cave colonization in North American cavefishes"

Ahrens, N., Elbers, D., Greb, H., Janssen-Bienhold, U., Koch, K.W. (2020) Interaction of G protein-coupled receptor kinases and recoverin isoforms is determined by localization in zebrafish photoreceptors. Biochimica et biophysica acta. Molecular cell research. 1868(4):118946.

Akira Ohkia, Youji Hu, Min Wang, Fernando U. Garcia, Mark E. Stearns; Evidence for Prostate Cancer-Associated Diagnostic Marker-1: Immunohistochemistry and in Situ Hybridization Studies. Clin Cancer Res 1 April 2004; 10 (7): 2452–2458. <https://doi.org/10.1158/1078-0432.CCR-03-0170>

Amsterdam A, Sadler KC, Lai K, Farrington S, Bronson RT, et al. (2004) Many Ribosomal Protein Genes Are Cancer Genes in Zebrafish. PLOS Biology 2(5): e139. <https://doi.org/10.1371/journal.pbio.0020139>

Ansar, M., Ebstein, F., Özkoç, H., Paracha, S.A., Iwaszkiewicz, J., Gesemann, M., Zoete, V., Ranza, E., Santoni, F.A., Sarwar, M.T., Ahmed, J., Krüger, E., Bachmann-Gagescu, R., Antonarakis, S.E. (2020) Biallelic variants in PSMB1 encoding the proteasome subunit β6 cause impairment of proteasome function, microcephaly, intellectual disability, developmental delay and short stature. Human molecular genetics. 29(7):1132-1143.

Ben-Moshe, Z., Vatine, G., Alon, S., Tovin, A., Mracek, P., Foulkes, N.S., and Gothilf, Y. (2010) Multiple PAR and E4BP4 bZIP transcription factors in zebrafish: diverse spatial and temporal expression patterns. Chronobiology International. 27(8):1509-1531.

Blanco-Sánchez, B., Clément, A., Fierro Jr, J., Washbourne, P., & Westerfield, M. (2014). Complexes of Usher proteins preassemble at the endoplasmic reticulum and are required for trafficking and ER homeostasis. Disease models & mechanisms, 7(5), 547-559.

Carney, T.J., Feitosa, N.M., Sonntag, C., Slanchev, K., Kluger, J., Kiyozumi, D., Gebauer, J.M., Coffin Talbot, J., Kimmel, C.B., Sekiguchi, K., Wagener, R., Schwarz, H., Ingham, P.W., and Hammerschmidt, M. (2010) Genetic analysis of fin development in zebrafish identifies furin and hemicentin1 as potential novel Fraser Syndrome disease genes. PLoS Genetics. 6(4):e1000907.

Chaudhari, S., Ware, A.P., Jayaram, P., Gorthi, S.P., El-Khamisy, S.F., Satyamoorthy, K. (2021) Apurinic/Apyrimidinic Endonuclease 2 (APE2): An ancillary enzyme for contextual base excision repair mechanisms to preserve genome stability. Biochimie. 190:70-90.

Chen, A., Singh, C., Oikonomou, G., & Prober, D. A. (2017). Genetic analysis of histamine signaling in larval zebrafish sleep. Eneuro, 4(1).

Cheng Chen, Haigen Huang, Ruibin Yan, Shuo Lin, Wei Qin, Loss of rps9 in Zebrafish Leads to p53-Dependent Anemia, G3 Genes|Genomes|Genetics, Volume 9, Issue 12, 1 December 2019, Pages 4149–4157, <https://doi.org/10.1534/g3.119.400585>

Chou, S.W., Hwang, P., Gomez, G., Fernando, C.A., West, M.C., Pollock, L.M., Lin-Jones, J., Burnside, B., and McDermott, B.M. (2011) Fascin 2b Is a Component of Stereocilia that Lengthens Actin-Based Protrusions. PLoS One. 6(4):e14807.

Cortés, R., Agulleiro, M.J., Navarro, S., Guillot, R., Sánchez, E., Cerdá-Reverter, J.M. (2014) Melanocortin receptor accessory protein 2 (MRAP2) interplays with the zebrafish melanocortin 1 receptor (MC1R) but has no effect on its pharmacological profile. General and comparative endocrinology. 201:30-6.

Coppola, U., Annona, G., D'Aniello, S., Ristoratore, F. (2016) Rab32 and Rab38 genes in chordate pigmentation: an evolutionary perspective. BMC Evolutionary Biology. 16:26.

Dooley, C.M., Schwarz, H., Mueller, K.P., Mongera, A., Konantz, M., Neuhauss, S.C., Nüsslein-Volhard, C., and Geisler, R. (2013) Slc45a2 and V-ATPase are regulators of melanosomal pH homeostasis in zebrafish, providing a mechanism for human pigment evolution and disease. Pigment cell & melanoma research. 26(2):205-217

Flores, M.V., Hall, C., Jury, A., Crosier, K., and Crosier, P. (2007) The zebrafish retinoid-related orphan receptor (ror) gene family. Gene expression patterns : GEP. 7(5):535-543.

Fries, R., Scholten, A., Säftel, W., and Koch, K.W. (2013) Zebrafish guanylate cyclase type 3 signaling in cone photoreceptors. PLoS One. 8(8):e69656.

Gaudet, P., Livstone, M., Thomas, P., The Reference Genome Project (2010) Annotation inferences using phylogenetic trees. Automated Data Submission. .

Gaudet, P., Livstone, M. S., Lewis, S. E., & Thomas, P. D. (2011). Phylogenetic-based propagation of functional annotations within the Gene Ontology consortium. Briefings in bioinformatics, 12(5), 449-462.

Gray, R. S., Wilm, T. P., Smith, J., Bagnat, M., Dale, R. M., Topczewski, J., ... & Solnica-Krezel, L. (2014). Loss of col8a1a function during zebrafish embryogenesis results in congenital vertebral malformations. Developmental biology, 386(1), 72-85.

Greiling, T. M., Houck, S. A., & Clark, J. I. (2009). The zebrafish lens proteome during development and aging. Molecular vision, 15, 2313.

Guillot, R., Muriach, B., Rocha, A., Rotllant, J., Kelsh, R.N., Cerdá-Reverter, J.M. (2016) Thyroid Hormones Regulate Zebrafish Melanogenesis in a Gender-Specific Manner. PLoS One. 11:e0166152.

Hartwig, C., Monis, W. J., Chen, X., Dickman, D. K., Pazour, G. J., & Faundez, V. (2018). Neurodevelopmental disease mechanisms, primary cilia, and endosomes converge on the BLOC‐1 and BORC complexes. Developmental neurobiology, 78(3), 311-330.

Heude, E., Shaikho, S., Ekker, M. (2014) The dlx5a/dlx6a Genes Play Essential Roles in the Early Development of Zebrafish Median Fin and Pectoral Structures. PLoS One. 9:e98505.

Jin, Z., Song, M., Wang, J., Zhu, W., Sun, D., Liu, H., & Shi, G. (2022). Integrative multiomics evaluation reveals the importance of pseudouridine synthases in hepatocellular carcinoma. Frontiers in genetics, 13, 944681

Khan, Z. A., Yumnamcha, T., Rajiv, C., Devi, H. S., Mondal, G., Devi, S. D., ... & Chattoraj, A. (2016). Melatonin biosynthesizing enzyme genes and clock genes in ovary and whole brain of zebrafish (Danio rerio): Differential expression and a possible interplay. General and Comparative Endocrinology, 233, 16-31.

Koc, E. C., Cimen, H., Kumcuoglu, B., Abu, N., Akpinar, G., Haque, M. E., ... & Koc, H. (2013). Identification and characterization of CHCHD1, AURKAIP1, and CRIF1 as new members of the mammalian mitochondrial ribosome. Frontiers in physiology, 4, 183.

Kuang, G.; Tao, W.; Zheng, S.; Wang, X.; Wang, D. Genome-Wide Identification, Evolution and Expression of the Complete Set of Cytoplasmic Ribosomal Protein Genes in Nile Tilapia. Int. J. Mol. Sci. 2020, 21, 1230. <https://doi.org/10.3390/ijms21041230>

Lavdovskaia, E., Denks, K., Nadler, F., Steube, E., Linden, A., Urlaub, H., ... & Richter-Dennerlein, R. (2020). Dual function of GTPBP6 in biogenesis and recycling of human mitochondrial ribosomes. Nucleic Acids Research, 48(22), 12929-12942.

Lim, S. K., & Gopalan, G. (2007). Aurora-A kinase interacting protein 1 (AURKAIP1) promotes Aurora-A degradation through an alternative ubiquitin-independent pathway. Biochemical Journal, 403(1), 119-127.

Lister, J., Close, J., and Raible, D. (2001) Duplicate mitf genes in zebrafish: complementary expression and conservation of melanogenic potential. Developmental Biology. 237(2):333-344.

Liu, F., Qin, Y., Huang, Y., Gao, P., Li, J., Yu, S., Jia, D., Chen, X., Lv, Y., Tu, J., Sun, K., Han, Y., Reilly, J., Shu, X., Lu, Q., Tang, Z., Xu, C., Luo, D., Liu, M. (2022) Rod genesis driven by mafba in an nrl knockout zebrafish model with altered photoreceptor composition and progressive retinal degeneration. PLoS Genetics. 18:e1009841.

Meserve, J. H., Nelson, J. C., Marsden, K. C., Hsu, J., Echeverry, F. A., Jain, R. A., ... & Granato, M. (2021). A forward genetic screen identifies Dolk as a regulator of startle magnitude through the potassium channel subunit Kv1. 1. PLoS genetics, 17(6), e1008943.

Nadauld, L.D., Chidester, S., Shelton, D.N., Rai, K., Broadbent, T., Sandoval, I.T., Peterson, P.W., Manos, E.J., Ireland, C.M., Yost, H.J., and Jones, D.A. (2006) Dual roles for adenomatous polyposis coli in regulating retinoic acid biosynthesis and Wnt during ocular development. Proceedings of the National Academy of Sciences of the United States of America. 103(36):13409-13414.

Nambiar, R.M., Ignatius, M.S., and Henion, P.D. (2007) Zebrafish colgate/hdac1 functions in the non-canonical Wnt pathway during axial extension and in Wnt-independent branchiomotor neuron migration. Mechanisms of Development. 124(9-10):682-698.

Oommen, S. K. (2006). Comparative analysis of human chromosome 22 CES-DGCR syntenic regions in chimpanzee, baboon, bovine, mouse and zebrafish and expression profiling in zebrafish early developmental stages using whole mount in situ hybridization. The University of Oklahoma.

Ogawa, Y., & Corbo, J. C. (2021). Partitioning of gene expression among zebrafish photoreceptor subtypes. Scientific reports, 11(1), 17340.

Pellacani, C., Bucciarelli, E., Renda, F., Hayward, D., Palena, A., Chen, J., ... & Somma, M. P. (2018). Splicing factors Sf3A2 and Prp31 have direct roles in mitotic chromosome segregation. Elife, 7, e40325.

Peng, X., Shang, G., Wang, W., Chen, X., Lou, Q., Zhai, G., Li, D., Du, Z., Ye, Y., Jin, X., He, J., Zhang, Y., Yin, Z. (2017) Fatty Acid Oxidation in Zebrafish Adipose Tissue Is Promoted by 1α,25(OH)_2_D_3_.. Cell Reports. 19:1444-1455.

Qian, M., Yao, S., Jing, L., He, J., Xiao, C., Zhang, T., Meng, W., Zhu, H., Xu, H., and Mo, X. (2013) ENC1-like Integrates the Retinoic Acid/FGF Signaling Pathways to Modulate Ciliogenesis of Kupffer's Vesicle during Zebrafish Embryonic Development. Developmental Biology. 374(1):85-95..

Rauch, G.J., Lyons, D.A., Middendorf, I., Friedlander, B., Arana, N., Reyes, T., and Talbot, W.S. (2003) Submission and Curation of Gene Expression Data. ZFIN Direct Data Submission. . (<http://zfin.org>).

Roosild, T. P., Castronovo, S., Villoso, A., Ziemba, A., & Pizzorno, G. (2011). A novel structural mechanism for redox regulation of uridine phosphorylase 2 activity. Journal of structural biology, 176(2), 229-237.

Shim, H., Kim, J. H., Kim, C. Y., Hwang, S., Kim, H., Yang, S., ... & Lee, I. (2016). Function-driven discovery of disease genes in zebrafish using an integrated genomics big data resource. Nucleic acids research, 44(20), 9611-9623.

Sun, P., Zhang, H., Shi, J., Xu, M., Cheng, T., Lu, B., ... & Huang, J. (2023). KRTCAP2 as an immunological and prognostic biomarker of hepatocellular carcinoma. Colloids and Surfaces B: Biointerfaces, 222, 113124.

Sun, C., Li, T., Song, X., Huang, L., Zang, Q., Xu, J., ... & Abliz, Z. (2019). Spatially resolved metabolomics to discover tumor-associated metabolic alterations. Proceedings of the National Academy of Sciences, 116(1), 52-57.

Sundvik, M., Kudo, H., Toivonen, P., Rozov, S., Chen, Y.C., and Panula, P. (2011) The histaminergic system regulates wakefulness and orexin/hypocretin neuron development via histamine receptor H1 in zebrafish. FASEB journal : official publication of the Federation of American Societies for Experimental Biology. 25(12):4338-47.

Sundvik, M., Chen, Y. C., & Panula, P. (2013). Presenilin1 regulates histamine neuron development and behavior in zebrafish, *Danio rerio*. Journal of Neuroscience, 33(4), 1589-1597.

Thisse, B., Thisse, C. (2004) Fast Release Clones: A High Throughput Expression Analysis. ZFIN Direct Data Submission. . (<http://zfin.org>).

Thisse, C., and Thisse, B. (2005) High Throughput Expression Analysis of ZF-Models Consortium Clones. ZFIN Direct Data Submission. . (<http://zfin.org>)

Thisse, C., and Thisse, B. (2008) Expression from: Unexpected Novel Relational Links Uncovered by Extensive Developmental Profiling of Nuclear Receptor Expression. ZFIN Direct Data Submission. . (<http://zfin.org>).

Tsai, S.M., Chu, K.C., Jiang, Y.J. (2020) Newly identified Gon4l/Udu-interacting proteins implicate novel functions. Scientific Reports. 10:14213.

Verreijdt, L., Debiais-Thibaud, M., Borday-Birraux, V., Van der Heyden, C., Sire, J.Y., and Huysseune, A. (2006) Expression of the dlx gene family during formation of the cranial bones in the zebrafish (Danio rerio): Differential involvement in the visceral skeleton and braincase. Developmental Dynamics : an official publication of the American Association of Anatomists. 235(5):1371-1389.

Wang, Z., Nishimura, Y., Shimada, Y., Umemoto, N., Hirano, M., Zang, L., Oka, T., Sakamoto, C., Kuroyanagi, J., and Tanaka, T. (2009) Zebrafish beta-adrenergic receptor mRNA expression and control of pigmentation. Gene. 446(1):18-27.

Xu, H., Ye, D., Behra, M., Burgess, S., Chen, S., and Lin, F. (2014) Gbeta1 controls collective cell migration by regulating the protrusive activity of leader cells in the posterior lateral line primordium. Developmental Biology. 385(2):316-27.

Yamaguchi, H., Oda, T., Kikkawa, M., Takeda, H. (2018) Systematic studies of all PIH proteins in zebrafish reveal their distinct roles in axonemal dynein assembly. eLIFE. 7.

Yu, S., Li, H., Gao, F., Zhou, Y. (2015) Crystal structure and potential physiological role of zebra fish thioesterase superfamily member 2 (fTHEM2). Biochemical and Biophysical Research Communications. 463:912-6.

Zhang, D., Xi, Y., Coccimiglio, M.L., Mennigen, J.A., Jonz, M., Ekker, M., and Trudeau, V.L. (2012) Functional prediction and physiological characterization of a novel short trans-membrane protein 1 (Stmp1) as a subunit of mitochondrial respiratory complexes. Physiological Genomics. 44(23):1133-1140.

Zhang, J., Jing, M., Li, P., Sun, L., Pi, X., Jiang, N., Zhu, K.K., Li, H., Li, J., Wang, M., Zhang, J., Liu, M., Mu, H., Hu, Y., Cui, X. (2023) Knockout of DLIC1 leads to retinal cone degeneration via disturbing Rab8 transport in zebrafish. Biochimica et biophysica acta. Molecular basis of disease. 1869(4):166645.

Zhou, Z., Peng, X., Chen, J., Wu, X., Wang, Y., Hong, Y. (2016) Identification of zebrafish magnetoreceptor and cryptochrome homologs. Science China. Life sciences. 59(12):1324-1331.

.
